# An efficient *in vivo*-inducible CRISPR interference system for group A *Streptococcus* genetic analysis and pathogenesis studies

**DOI:** 10.1101/2024.02.22.581527

**Authors:** Elisabet Bjånes, Alexandra Stream, Axel B. Janssen, Paddy S. Gibson, Afonso M. Bravo, Samira Dahesh, Jonathon L. Baker, Andrew Varble, Victor Nizet, Jan-Willem Veening

**Author notes:** Contributed equally (Elisabet Bjånes, Alexandra Stream). Correspondence to Jan-Willem Veening, or Victor Nizet.

## Abstract

While genome-wide transposon mutagenesis screens have identified numerous essential genes in the significant human pathogen *Streptococcus pyogenes* (group A *Streptococcus* or GAS), many of their functions remain elusive. This knowledge gap is attributed in part to the limited molecular toolbox for controlling GAS gene expression and the bacterium’s poor genetic transformability. CRISPR interference (CRISPRi), using catalytically inactive GAS Cas9 (dCas9), is a powerful approach to specifically repress gene expression in both bacteria and eukaryotes, but ironically has never been harnessed for controlled gene expression in GAS. In this study, we present a highly transformable and fully virulent serotype M1T1 GAS strain and introduce a doxycycline-inducible CRISPRi system for efficient repression of bacterial gene expression. We demonstrate highly efficient, oligo-based sgRNA cloning directly to GAS, enabling the construction of a gene knockdown strain in just two days, in contrast to the several weeks typically required. The system is shown to be titratable and functional both *in vitro* and *in vivo* using a murine model of GAS infection. Furthermore, we provide direct *in vivo* evidence that the expression of the conserved cell division gene *ftsZ* is essential for GAS virulence, highlighting its promise as a target for emerging FtsZ-inhibitors. Finally, we introduce SpyBrowse (https://veeninglab.com/SpyBrowse), a comprehensive and user-friendly online resource for visually inspecting and exploring GAS genetic features. The tools and methodologies described in this work are poised to facilitate fundamental research in GAS, contribute to vaccine development, and aid in the discovery of antibiotic targets.

**Significance statement:** While GAS remains a predominant cause of bacterial infections worldwide, there are limited genetic tools available to study its basic cell biology. Here, we bridge this gap by creating a highly transformable, fully virulent M1T1 GAS strain. In addition, we established a tight and titratable doxycycline-inducible system and developed CRISPR interference for controlled gene expression in GAS. We show that CRISPRi is functional *in vivo* in a mouse infection model. Additionally, we present SpyBrowse, an intuitive and accessible genome browser (https://veeninglab.com/SpyBrowse). Overall, this work overcomes significant technical challenges of working with GAS, and together with SpyBrowse, represents a valuable resource for researchers in the GAS field.

## Introduction

*Streptococcus pyogenes*, also known as group A *Streptococcus* (GAS), is a bacterium commonly present in the throat and on the skin (1, 2). This pathogen is notorious for causing strep throat and impetigo, accounting for approximately 700 million non-invasive infections each year (3–5). However, GAS can also lead to serious invasive diseases, including necrotizing fasciitis and streptococcal toxic shock syndrome, resulting in over 150,000 deaths annually (4). Additionally, GAS is the immunological trigger for acute rheumatic fever (ARF) and rheumatic heart disease (RHD), causing substantial death and disability in many developing countries. Despite rising GAS resistance to certain antibiotic classes, the pathogen has fortunately remained susceptible to penicillin and other β-lactam agents (6).

There is presently no commercially available vaccine to protect against GAS infection (7). GAS presents a challenge for vaccine antigen selection due to the variability in the abundant surface-exposed M protein with over 230 *emm* types circulating globally (8). The most common *emm* type, M1, is a major contributor to GAS global epidemiology, and is particularly prominent in severe, invasive infections (9). The search for new GAS antibiotic targets and vaccine candidates is hindered by a knowledge gap in fundamental GAS biology, partly because M1-type GAS strains are exceptionally challenging to manipulate genetically (10, 11). In this study, we present a toolbox for GAS genetic engineering, utilizing the hard-to-transform and clinically relevant M1T1-type strain 5448 (NV1) as model (1, 12). We selected strain 5448 since it is commonly used, and we reckoned that if our approaches work in this strain, they are highly likely to also work in generally easier to work with GAS strains. This toolbox should be generally applicable to GAS and related bacteria, encompassing protocols for recombineering using GoldenGate-assembled linear DNA, oligo-based sgRNA cloning, a titratable doxycycline-inducible promoter, and CRISPR interference (CRISPRi) effective both *in vitro* and *in vivo* in a murine GAS infection model.

Additionally, we present SpyBrowse, an intuitive and accessible genome browser (https://veeninglab.com/SpyBrowse), based on JBrowse 2 (13), a graphical and user-friendly interface to explore the GAS genomic landscape with direct linking to UniProt (14) and other useful resources such as PaperBlast (15). Overall, this work overcomes significant technical challenges of working with GAS, facilitating genetic engineering and targeted gene knockdowns to advance our insights into the physiology and cell biology of this preeminent human bacterial pathogen.

## Results

### Improved transformability of GAS M1T1 Strain 5448 by mutating *hsdR*

GAS5448, a widely used strain in fundamental research, serves as a clinical representative of the globally distributed M1T1 serotype associated with severe invasive infections. While 5448 has been effectively employed in murine models of GAS infection (16, 17), its genetic manipulation poses challenges, with even the construction of transposon mutant libraries proving highly difficult (10, 11, 18). To enhance GAS 5448 transformation efficiencies while retaining full virulence, we targeted one of the major barriers to transformation—the HsdR restriction subunit of the conserved three-component Type I restriction-modification (RM) system, HsdRSM. *Hsd,* denoting <u>H</u>ost <u>s</u>pecificity of <u>D</u>NA, signifies how these Type I RM systems cleave intracellular (foreign) DNA with improper methylation patterns. Mutations in this system improve transformation efficiency in other GAS strains (19–22), but with potential pleiotropic consequences. For example, while deletion of the entire *hsdRSM* system in serotype M28 GAS strain MEW123 boosted transformation efficiency, it concurrently reduced virulence in a murine model of infection (20). A spectinomycin marker-replacement mutant eliminating just the restriction subunit *hsdR* also increased transformation efficiency but led to partially methylated genomic DNA likely due to polar effects (20).

To address the above obstacles, we generated a mutant of *hsdR* in the wild-type (WT) GAS 5448 (strain NV1, Table 1) using an erythromycin resistance cassette. This cassette, previously demonstrated to provide selectable resistance at very low transcription levels without requiring an upstream promoter (23), was introduced using GoldenGate assembly combined with recombineering (Fig. 1A-B, strain NV28: *hsdR::ery*, see Methods for details). SMRT sequencing (PacBio) confirmed the desired mutation in NV28, and the identical genomic DNA methylation pattern to NV1 validating the absence of polar effects from the erythromycin cassette replacement. Global DNA methylation patterns analysis suggested that the MTase activity of the HsdRSM system of GAS strain 5448 methylates adenines at position 3 in the motifs 5’-GCANNNNNNTTAA’3’ and 5’-TTAANNNNNNTGC’3’, with all 344 motifs in the NV1 genome methylated, consistent with patterns observed in other M1 strains (21) (Fig. 1C).

**Fig. 1:**
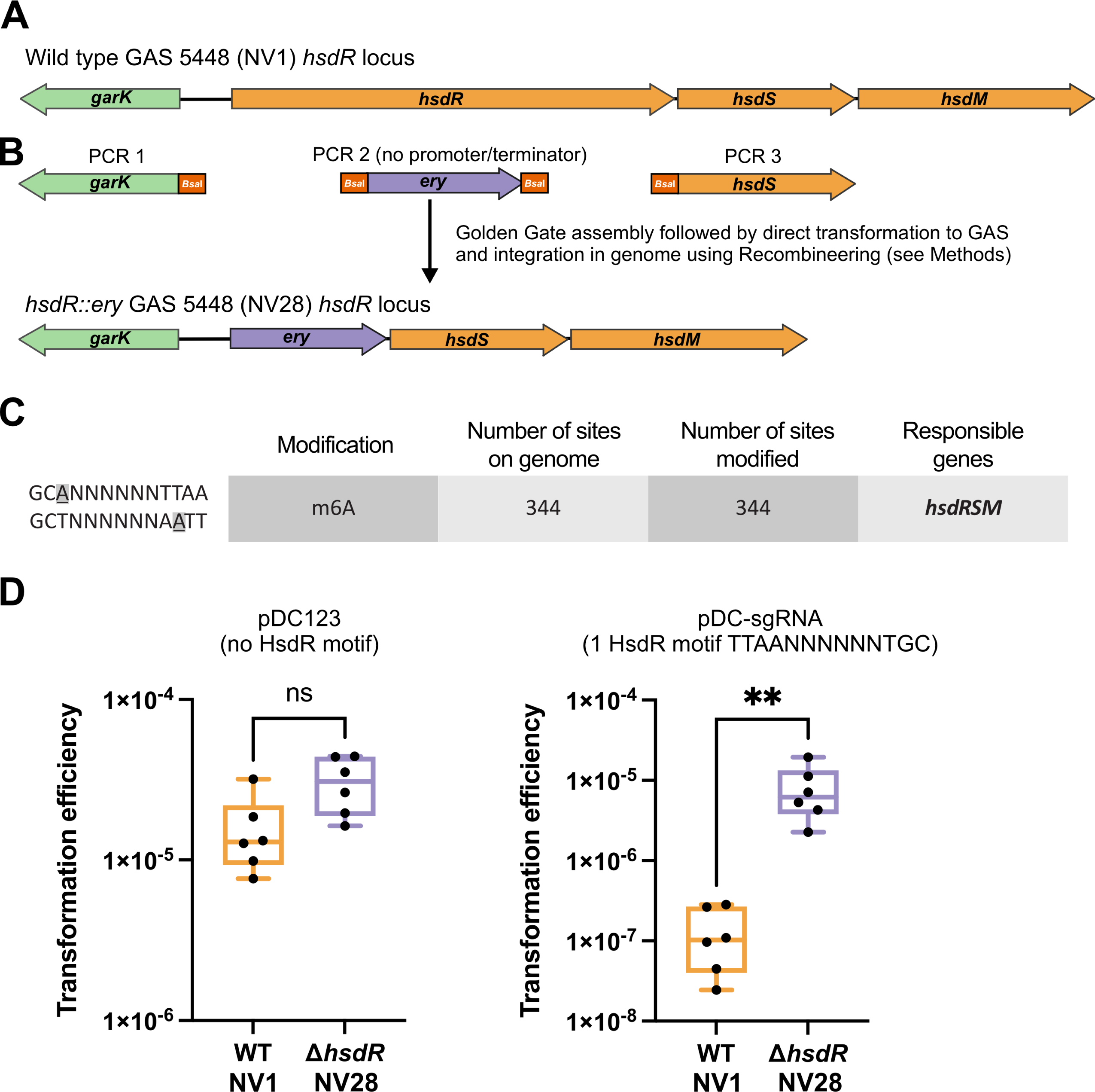
Golden Gate assembly and recombineering to create a highly transformable *hsdR* mutant in M1T1 GAS strain 5448. (**A**) Schematic representation of the genomic organization of the *hsdRSM* locus in WT GAS strain M1 5448 (NV1). Gene annotations were made by Prokka (43). (**B**) Three PCRs were performed each containing unique *Bsa*I restriction sites that allow for scarless Golden Gate assembly of a linear fragment in which *hsdR* is replaced by an erythromycin resistance cassette (*ery*). PCR 1 contains 1392 bp upstream of *hsdR* (including the *garK* gene and excluding the start codon of *hsdR*), PCR 2 contains the 785 bp promoterless and terminatorless *ery* cassette (23) flanked by *Bsa*I sites while PCR 3 contains a 1238 bp downstream region of *hsdR* (including its stop codon and *hsdS*). After Golden Gate assembly with *Bsa*I and T4 ligase (see Methods), the resulting 3383 bp ligation product was purified and directly used to transform electrocompetent NV2 (NV1 + pAV488) cells that were grown with 1 mM of IPTG to induce the recombineering system present on plasmid pAV488 resulting in strain NV3 (NV2, *hsdR::ery*). Recombineering of GAS using pAV488 is described in more detail elsewhere (Varble, unpublished). Strain NV3 was plasmid cured resulting in strain NV28 (NV1, *hsdR::ery*). The genomic organization of the *hsdRSM* locus in strain NV28 (*hsdR::ery*) is shown. (**C**) Methylation motifs identified in strains NV1, NV28, and NV6 using SMRT sequencing (see Methods for details). (**D**) Transformation efficiencies of WT GAS 5448 (NV1) and the *hsdR::ery* mutant (NV28) are shown. Left: Strains were transformed with 140 ng (∼50 fmol) of plasmid pDC123 (24) that does not contain an HsdR-motif. Right: Strains were transformed with 247 ng (∼95 fmol) of plasmid pDC-sgRNA (see Fig. 2C) that contains a single HsdR-motif. Each dot represents a replicate and a Kolmogorov-Smirnov test was used to calculate statistical significant differences between the two strains (P value **<0.005).

**Table 1:**
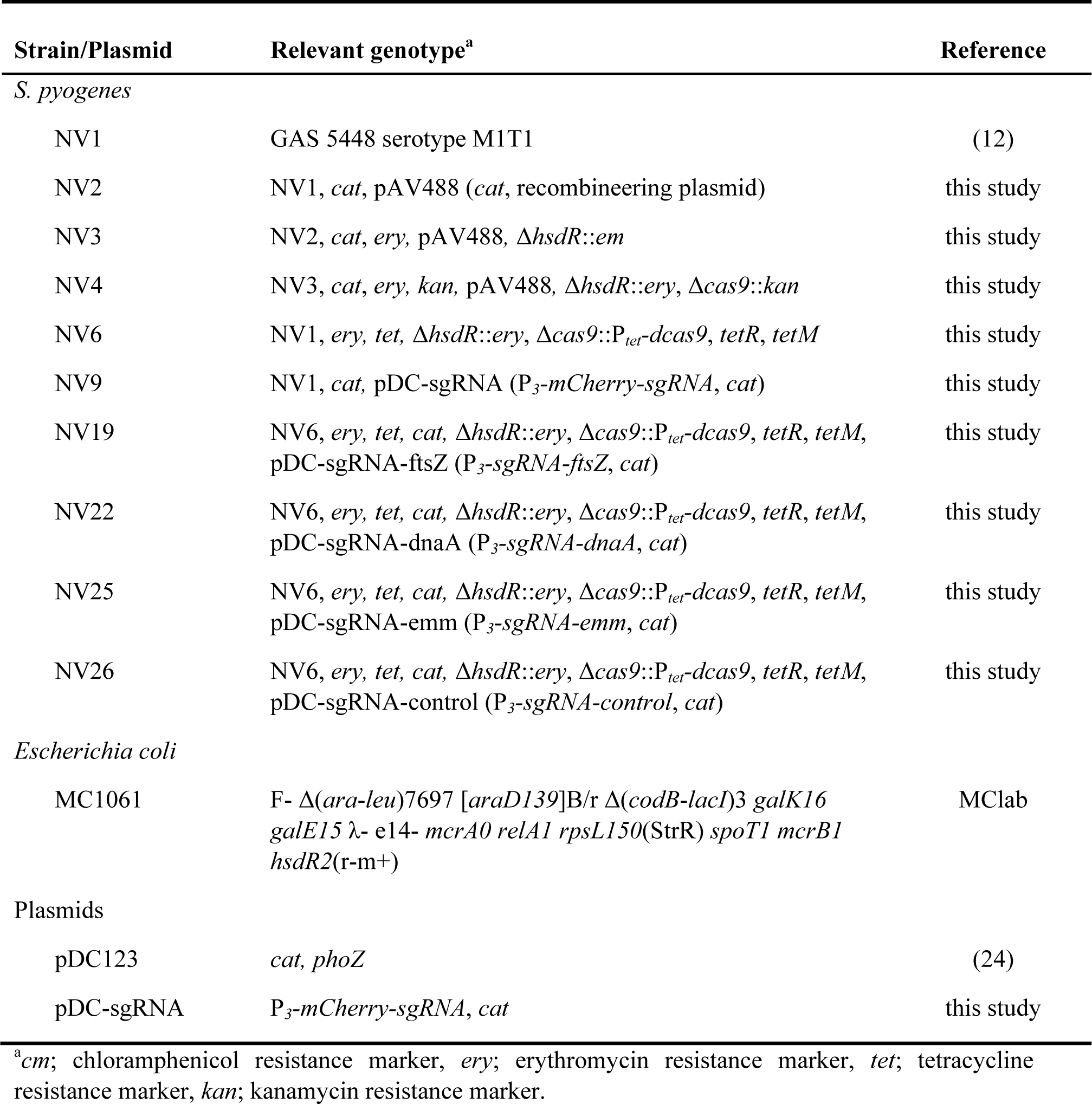
Plasmids and bacterial strains used in this study.

Electroporation transformation efficiency with replicative plasmid pDC123 (24), lacking a putative HsdR restriction site, remained as efficient in the *hsdR::ery* mutant as in the WT. However, when strain NV28 was transformed with replicative plasmid pDC-sgRNA (see below, Fig. 2E), containing a single HsdR site, transformation efficiencies were more than 60-fold higher than transforming pDC-sgRNA to WT NV1 (Fig. 1D). Consequently, the newly constructed GAS M1T1 5448-derived *hsdR::ery* mutant not only allows for increased transformation efficiencies with foreign DNA containing HsdR motifs, but may also serve as a valuable genetic background for future studies. Moreover, the virulence of the strain harboring the *hsdR::ery* cassette was indistinguishable from its WT parent strain (see below, Fig. 5C).

**Fig. 2:**
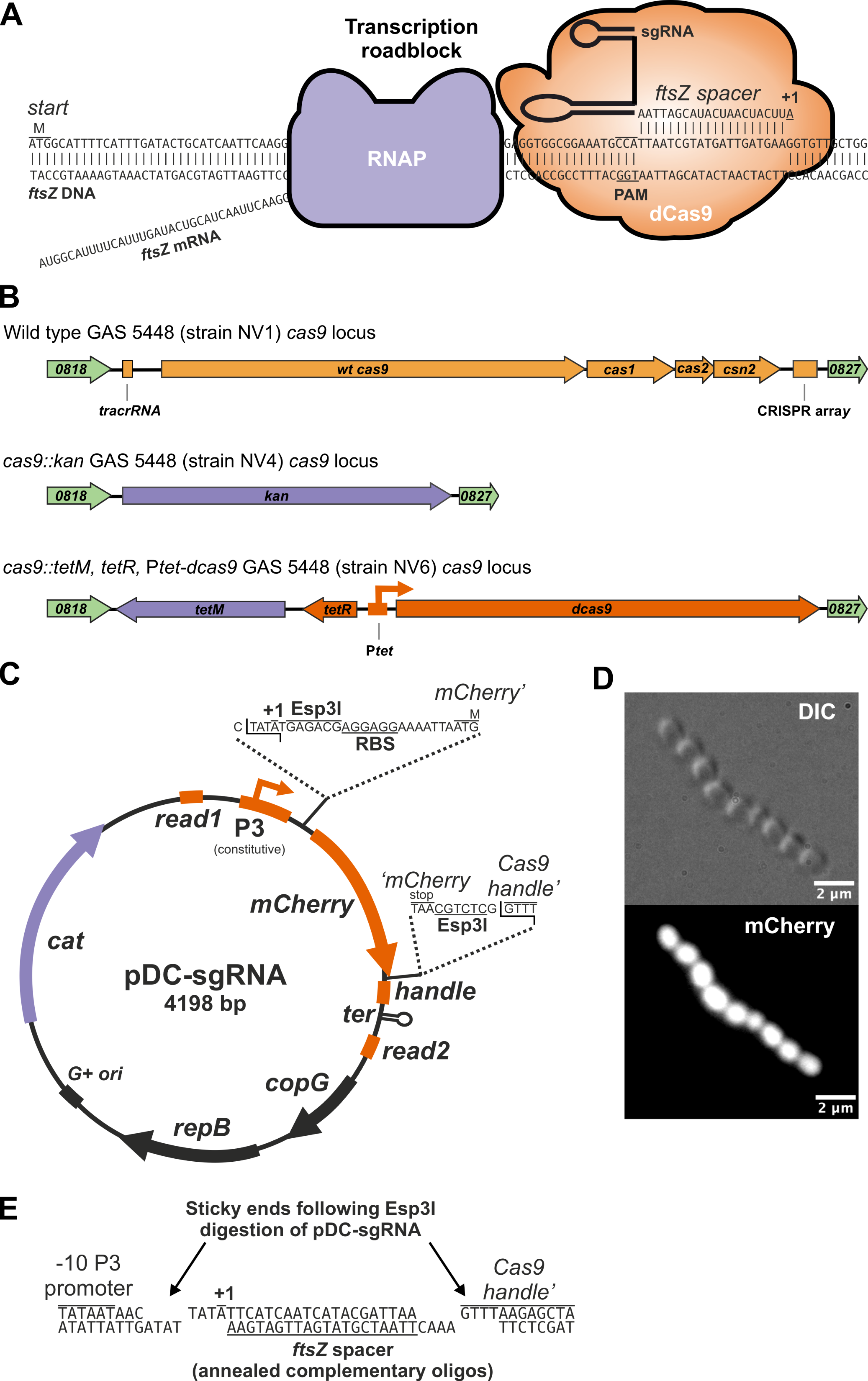
Development of a doxycycline-inducible CRISPRi system for use in Group A Streptococcus. (**A**) Schematic representation of CRISPRi. An sgRNA encoding a 20 nt spacer region targets dCas9 to the non-template strand containing a protospacer adjacent motif (PAM) on the GAS *ftsZ* locus. Transcription elongation by RNA polymerase (RNAP) is consequently hampered leading to reduced levels of FtsZ in the cell when dCas9 is induced. (**B**) Schematic representation of the genomic organization of the *cas9* locus in wildtype GAS strain 5448 (NV1), in strain NV4 (*cas9::kan*) and strain NV6 (*cas9::tetM*, *tetR*, Ptet-*dcas9*). (**C**) Schematic representation of sgRNA cloning vector pDC-sgRNA. (**D**) Strain NV9 (NV1, pDC-sgRNA) was grown in THB containing 2 μg/ml of chloramphenicol at 37°C and exponentially growing cells were imaged by differential interference contrast (DIC) and fluorescence microscopy. Scale bar: 2 μm. (**E**) Oligo-based sgRNA cloning in pDC-sgRNA. Plasmid pDC-sgRNA is cut with Esp3I (or its isoschizomer BsmBI) and its resulting sticky ends are shown. Two complementary oligos of each 24 nt long that include a 20 bp spacer sequence are annealed, phosphorylated, ligated and electroporated to competent GAS (see Methods). As an example, the two oligos used to clone the sgRNA targeting GAS *ftsZ* is shown (for oligo design, see Table S1). Successful clones lost the *mCherry* cassette and will be white on plate instead of pink.

The long-read genome-sequenced NV1 strain was compared to the published 5448 reference genome which was generated by illumina short read sequencing (25), revealing a large inversion of 1,475,033 base pairs. This structural variant is flanked by two ISAs1-like element IS1548 family transposase genes, which typically harbor terminal inverted repeats (26). The two transposase genes are oriented in opposing directions and exhibit two single nucleotide polymorphisms. It is highly probable that the short read approach used in the original 5448 genome project contributed to an inaccurate genome assembly. In addition, NV1 contains 6323 additional bases compared to 5448 and several SNPs mainly attributed to the transposase genes (see Methods). Therefore, we propose using the genome assembly presented here for GAS studies involving strain 5448 NV1 (Genbank accession CP140117.2).

### Replacing wild type GAS *cas9* with a doxycycline-inducible dead *cas9* (*dcas9*)

CRISPRi has emerged as a powerful approach for evaluating the functionality of both essential and non-essential genes across a wide range of bacteria (27–30). This gene-silencing technique employs a catalytically inactive version of the GAS Cas9 protein (dCas9) and a single guide RNA (sgRNA). The dCas9-sgRNA complex binds via complementary base pairing of the spacer sequence in the sgRNA to a specific genomic DNA sequence located beside a protospacer adjacent motif (PAM). This binding effectively halts transcription of the target gene and genes within the same operon (**Fig. 2A**) (31, 32). Curiously, while GAS Cas9 is frequently utilized in CRISPRi across various bacteria, as far as we are aware, CRISPRi has not been applied to GAS (33). Our objective in this study is to close this gap and introduce CRISPRi as a valuable tool for investigating gene function in GAS.

An optimal CRISPRi system should tightly regulate the expression of a dCas9. However, the selection of suitable inducible promoters for GAS has been limited. The most commonly used system involves a tetracycline-inducible promoter adapted from the Gram-negative bacterium *Escherichia coli* (34, 35). Unfortunately, this system has been reported to exhibit either leakiness or a narrow induction range in GAS (35). In a previous study, we developed a tetracycline, anhydrotetracycline, and doxycycline-inducible CRISPRi system with a highly dynamic range in *S. pneumoniae.* This was achieved by codon optimizing *E. coli tetR,* combined with the selection and counterselection of random promoter libraries containing *tetO* operators (23, 36). Given the similar codon usage between GAS and *S. pneumoniae*, we hypothesized that this TetR-based system could also function effectively in GAS. An additional advantage of this TetR-based system is the excellent tissue penetration of doxycycline (doxy) (37). Our prior work demonstrated the successful delivery of doxy through mouse chow or direct intraperitoneal injection to induce pneumococcal constructs in various mouse body/tissue niches, including the blood and lungs (23, 36, 38).

To mitigate potential crosstalk with the intrinsic Cas9-CRISPR system, we strategically chose the WT GAS *cas9* locus as the genomic integration site for stable introduction of the inducible *dcas9* cassette. Using Golden Gate assembly in conjunction with recombineering (see Methods), we first replaced the native *cas9* locus, inclusive of the tracrRNA and the CRISPR spacer array locus, with a kanamycin resistance marker (*kan*), resulting in strain NV4 (**Fig. 2B**). Next, we replaced the *kan* marker of strain NV4 with the *tetM*-*tetR*-P*tet-dcas9* cassette from *S. pneumoniae* strain VL3469 (36). This cassette imparts tetracycline/doxycycline resistance via the *tetM* marker and positions *dcas9* under the TetR-controlled *S. pneumoniae* P*tet* promoter (23). Finally, the strain underwent curing of the pAV488 recombineering plasmid, yielding strain NV6 (**Fig. 2B**). Whole-genome sequencing of strain NV6 verified the accurate introduction of all elements (SRA genome accession number SRX22828772).

### Construction of Gram-positive replicative vector pDC-sgRNA enables efficient sgRNA cloning

Next, we designed a replicative vector tailored for direct sgRNA cloning in GAS. We replaced the *phoZ* reporter gene within the Gram-positive high-copy number (∼24 - 90 copies per cell) rolling-circle replication pDC123 plasmid (24, 39) with an *mCherry* cassette flanked by the sgRNA scaffold sequence and Esp3I restriction sites. This configuration allows for Golden Gate assembly or direct ligation of sgRNA spacers using annealed oligonucleotides (**Fig. 2C**). This vector is also capable of replicating in *recA+ E. coli* strains such as MC1061, albeit at low copy numbers (∼4 copies per cell)(39). In addition, we incorporated Illumina read 1 and read 2 sequences flanking the sgRNA scaffold to facilitate CRISPRi-seq (40). Notably, we included the +1 of the P3 promoter, ensuring that all cloned spacers initiate with an adenine nucleotide (40). This design promotes efficient transcription and prevents undesirable uridines at the 5’ end of the sgRNA (41). Finally, the vector contains a chloramphenicol resistance cassette (*cat*) for selection in GAS (**Fig. 2C**). To validate the functionality of the synthetic P3 promoter (42) in GAS, we transformed pDC-sgRNA into strain NV1, creating NV9. Bacterial examination by fluorescence microscopy revealed strong and uniform mCherry expression in all cells (**Fig. 2D**), confirming the functionality of the P3 promoter in GAS. Given that pDC-sgRNA features the Gram-positive pLS1/pJS3 origin, it is anticipated to be functional across a broad range of Gram-positive bacteria, including but not limited to *Lactococcus lactis*, *Bacillus subtilis*, *S. pneumoniae*, *Enterococcus faecalis* and group B *Streptococcus*.

### An established pipeline generates unique genome-wide sgRNA spacers for GAS 5448

With the establishment of an inducible *dcas9* strain and an sgRNA expression system for GAS, we systematically designed sgRNA spacer sequences targeting every annotated genetic feature of the GAS 5448 genome using our previously established pipeline (40). The algorithm employed for spacer design only designs spacers that exclusively target the non-template strand (**Fig. 2A**) and considers factors such as specificity (to limit off target effects), distance to the start site, and avoiding Esp3I/BsmBI restriction sites (40). Upon running the pipeline, a total of 1823 unique spacers were generated, designed to target 1879 out of 1894 features on the NV1 (5448) genome as annotated by Prokka (43). Suitable spacers could not be identified for 15 genetic elements. Note that certain designed sgRNAs may target more than one genetic element if that element is present in multiple copies (e.g., repeats, gene duplications, etc.). A comprehensive list of all designed sgRNAs is provided in Table S1.

### SpyBrowse: *Streptococcus pyogenes* (GAS) genome browser

On the basis of the fully closed GAS NV1 genome (Genbank accession CP140117.2), in conjunction with automated genome annotation (see Methods), we have developed a new user-friendly genome browser named SpyBrowse, accessible at https://veeninglab.com/SpyBrowse. Based on JBrowse 2 (13), SpyBrowse empowers users to conveniently and flexibly search for their gene of interest using either its gene name or locus tag (**Fig. 3A**). Clicking on features reveals further information (**Fig. 3B**), including links to Uniprot (including subcellular location predictions and Alphafold protein structure predictions; **Fig. 3C**), NCBI protein entry (**Fig. 3D**), and PaperBlast (**Fig. 3E**). In addition, SpyBrowse incorporates a track displaying all designed sgRNAs from Table S1.

**Fig. 3:**
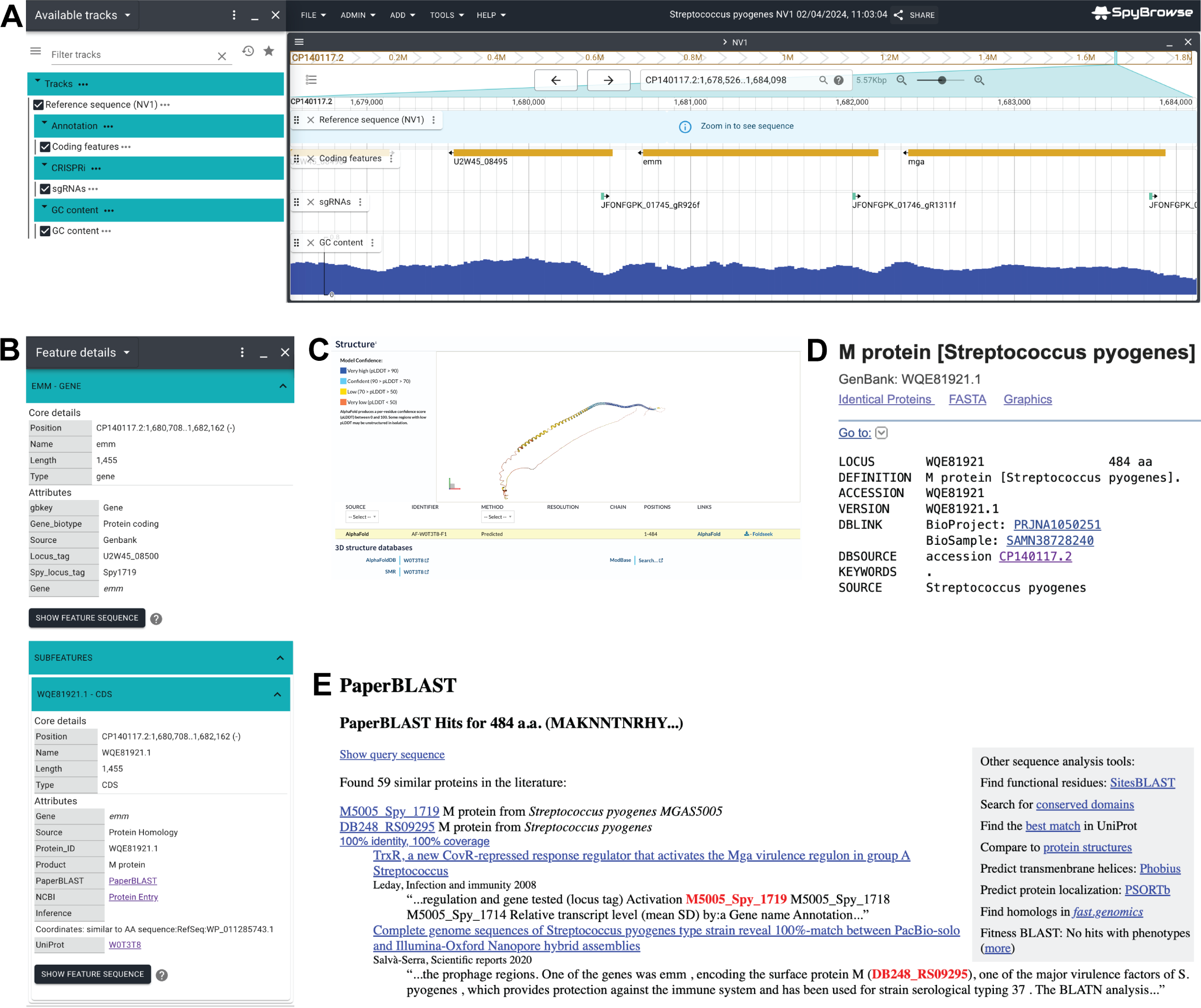
Spybrowse available at https://veeninglab.com/SpyBrowse. (**A**) A screenshot of the *emm* locus as shown in SpyBrowse. In the left pane, tracks can be turn on/off. In the right pane, the genome can be browsed via dragging the mouse to the left or right, zooming in and out, or searched on gene name and/or locus tags (e.g. Spy1719 or U2W45_08500 for *emm*). Annotated features such as genes, rRNAs, and tRNAs are displayed. The designed spacer targeting the non-template strand of *emm* (JFONFGPK_01746_gR1311f) is shown via the sgRNA track. (**B**) For each coding sequence, a context menu provides links to external resources, such as Uniprot (with Alphafold prediction), NCBI (with Blast function) and PaperBlast. (**C**) The predicted Alphafold structure of the M1 protein through the Uniprot link in SpyBrowse is shown. (**D**) The NCBI entry for the M1 protein is shown. (**E**) The Paperblast hits for the M1 protein are shown.

### CRISPRi can be used to tunably repress GAS gene expression

To assess the effectiveness of CRISPRi in downregulating the expression of essential genes in GAS, we targeted *dnaA*, which encodes the conserved replication initiator in bacteria, and *ftsZ*, responsible for the conserved tubulin-like cell division protein essential for cell division. FtsZ is a component of the highly conserved division and cell wall (*dcw*) genomic cluster (44). Due to anticipated polar effects of CRISPRi, it is expected that targeting *ftsZ* will also impact transcription of other members of the *dcw* cluster. To construct each sgRNA plasmid, the design outlined in Table S1 was used, and complementary oligos were synthesized for each target. These oligos were annealed, phosphorylated, and ligated in Esp3I-digested pDG-sgRNA (**Figs. 2C, 2E**). Additionally, a control sgRNA was included, containing a random 20-bp sequence (5’-CATACAAGTCGATAGAAGAT-3’) that does not match any region in the GAS NV1 5448 genome. The ligation mixtures were directly used to transform electrocompetent GAS NV6 cells (see Methods). Correct clones were cultured in Todd Hewitt broth (THB) within microtiter plates, and the optical density was measured every 10 min. As depicted in **Fig. 4A**, induction of dCas9 by increasing concentrations of doxycycline (doxy) resulted in reduced growth with the *dnaA* and *ftsZ* sgRNAs. but not with the control sgRNA. Note that at concentrations of 50 ng/ml of doxy, a reduction in growth is also observed for the control strain.

**Fig. 4:**
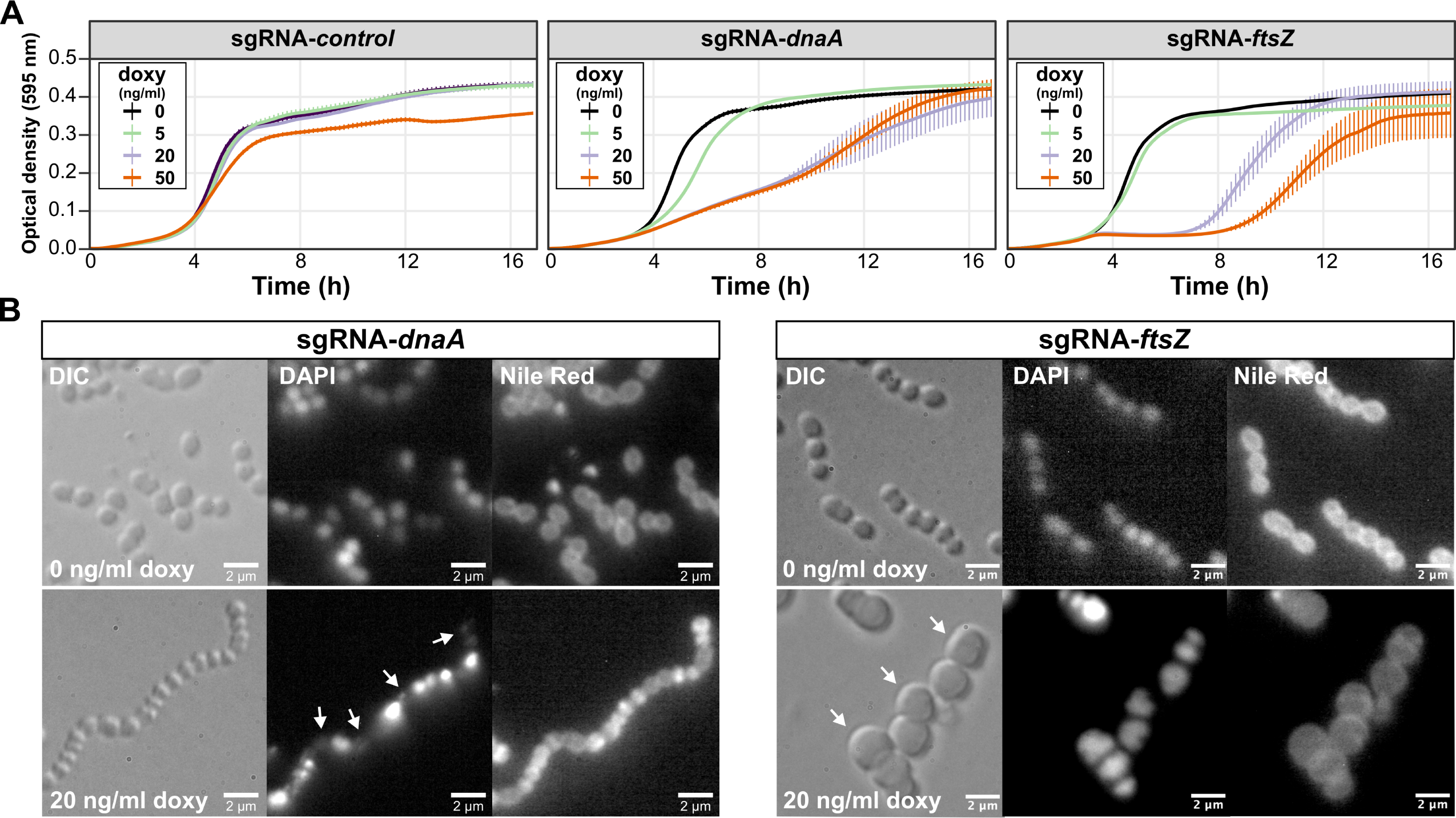
Targeted repression of *dnaA* and *ftsZ* gene expression by CRISPRi in GAS. (**A**) Strains NV26 (NV6+pDC-sgRNA-*control*), NV22 (NV6+pDC-sgRNA-*dnaA*) and NV19 (NV6+pDC-sgRNA-*ftsZ*) were grown in THB containing 2 μg/ml of chloramphenicol at 37°C. Exponentially growing cells were diluted to a start OD of 0.004 in a microtiter plate containing fresh THB with varying concentrations of doxycycline (doxy). The optical density at 595 nm was measured every 10 min. Each line is an average of three replicates with the standard deviation shown. For clarity, the optical density is plotted on a linear scale. (**B**) Strains NV22 and NV19 were grown in the presence or absence of 20 ng/ml of doxy and after 3h, cells were imaged by fluorescence microscopy. DAPI was used to stain the nucleoids and Nile red to stain the membrane. Scale bar: 2 μm. Arrows point to anucleate cells (sgRNA-*dnaA*) or cells with a block in division (sgRNA-*ftsZ*).

To confirm the specific targeting of *dnaA* and *ftsZ* by the designed sgRNAs, cells were grown in the presence or absence of 20 ng/ml of doxy for 3h and imaged by microscopy. As anticipated, repression of *dnaA* expression by CRISPRi resulted in cell chaining, with numerous anucleate cells indicative of a failure to initiate DNA replication (**Fig. 4B**). Targeting *ftsZ* led to a distinct block in cell division, resulting in enlarged cells (**Fig. 4B**). These findings collectively show that the CRISPRi system developed for GAS in this study is titratable, specific, and suitable for studying essential genes.

### CRISPRi to deplete expression of the M protein, a signature GAS virulence factor

To explore the applicability of the CRISPRi system described here for GAS in pathogenesis studies, we initially designed an sgRNA targeting the *emm* gene, encoding the M-protein. As shown in Fig. 5A, the growth of strain NV25 (NV6+pDC-sgRNA-*emm*) was not perturbed by the induction of dCas9 with doxy *in vitro*. Flow cytometry analysis, using an antibody specific to the M1T1 protein, revealed that NV25 induced with doxy exhibited approximately eight-fold less surface-exposed M-protein. This result underscores the precision and efficacy of the CRISPRi-based gene repression system developed for GAS (**Fig. 5B**).

**Fig. 5:**
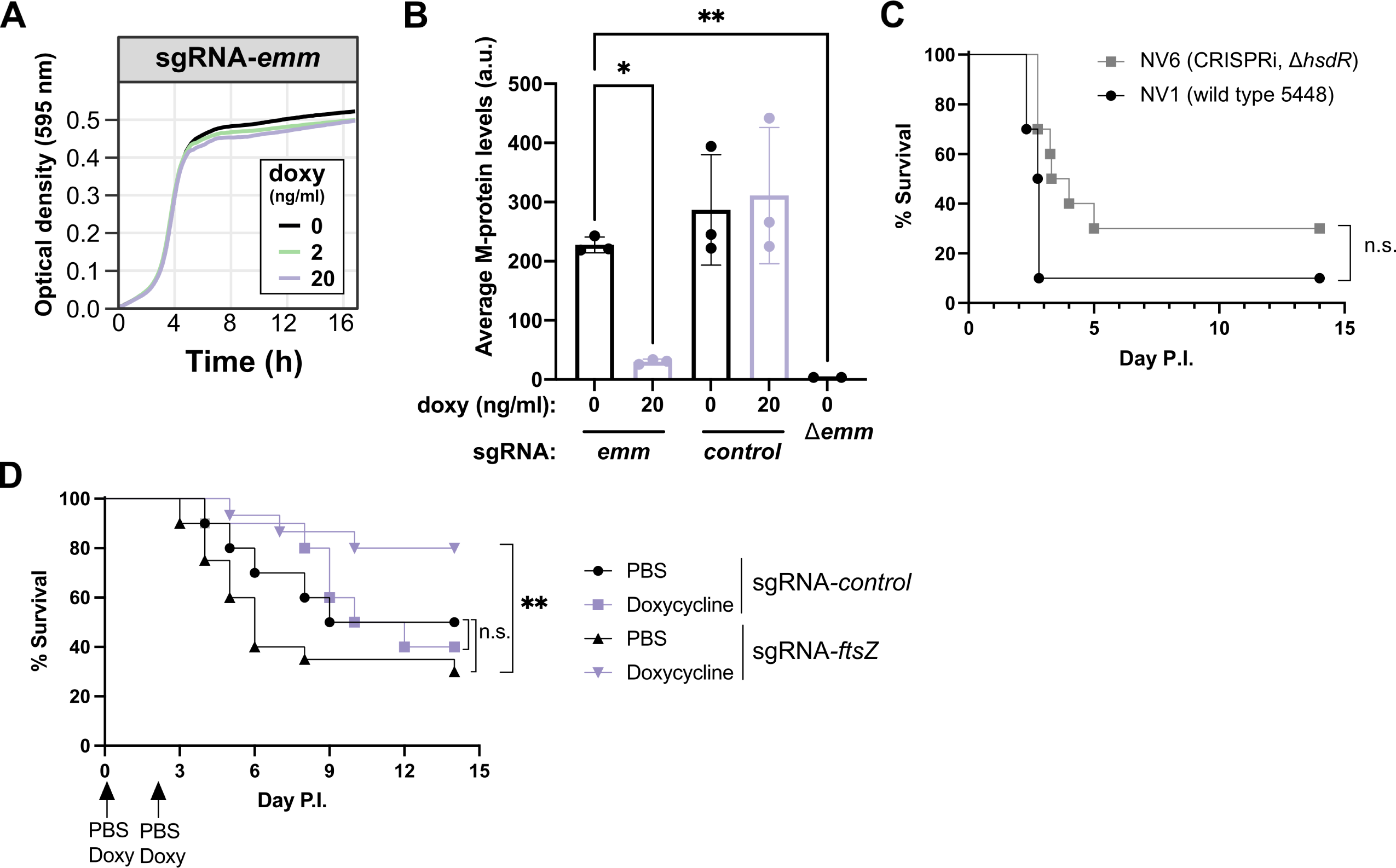
Efficient depletion of M-protein and *in vivo* CRISPRi in GAS. (**A**) Strain NV25 (NV6+pDC-sgRNA-*emm*) was grown in THB containing 2 μg/ml of chloramphenicol at 37°C in microtiter plates in the presence of various concentrations of doxycycline. (**B**) Strains NV25 (sgRNA-emm), NV26 (sgRNA-control), and M protein knockout GAS M1T1 5448 were grown in THB to mid-logarithmic growth in the presence or absence of 20 ng/ml doxycycline. Bacteria were immunostained using M protein antisera and analyzed by flow cytometry as described in the Methods. (**C**) Strains NV1 (wild type GAS 5448) and NV6 (*cas9::tetM*, *tetR*, Ptet-*dcas9*, *hsdR::ery*) were grown in THB at 37°C and 1-3*10^8^ CFU were used to infect CD-1 mice IP (10 mice per group). Disease score and survival was followed for 14 days. There was no statistical significant (n.s.) difference in virulence between both strains (Mantel-Cox Log Rank test). (**D**) Strains NV26 (NV6+pDC-sgRNA-*control*) and NV19 (NV6+pDC-sgRNA-*ftsZ*) were grown in THB containing 2 μg/ml of chloramphenicol at 37°C and 1-3*10^8^ CFU were used to infect CD-1 mice IP (10 mice per group). Disease score and survival was followed for 14 days. Doxy-induced mice infected with NV19 showed a statistically significant increased survival (** p = 0.0027, Mantel-Cox Log Rank test).

### CRISPRi targeting of *ftsZ* reveals contribution to *in vivo* GAS virulence

To assess whether the engineered GAS CRISPRi strain NV6 exhibits a comparable virulence profile to the WT 5448 NV1, we intraperitoneally (I.P.) infected outbred CD-1 mice with 1-3 x 10^8^ colony-forming units (CFU) and monitored disease progression. As shown in Fig. 5C, no significant difference in mouse survival was observed, demonstrating that the replacement of the CRISPR locus with our CRISPRi cassette, as well as the *hsdR* mutation, had no discernable global impact on systemic virulence.

A notable strength of CRISPRi lies in its ability to assess the functionality of essential genes. FtsZ, a highly conserved, tubulin-like, essential cell division protein, represents an attractive target for an expanding list of antibiotic candidates (38, 45). Certain compounds, such as PC190723, are particularly intriguing as they exhibit potent FtsZ inhibitory activity without targeting eukaryotic tubulin (46). While *in vivo* studies have demonstrated the efficacy of several FtsZ inhibitors in clearing bacterial infections, and it is evident that *ftsZ* is essential *in vitro* (**Fig. 4**), direct confirmation of the essentiality of FtsZ or the *dcw* cluster has not been established, to the best of our knowledge. CRISPRi was used to target *ftsZ* (and because of the polarity of CRISPRi, potentially other genes in the *dcw* cluster) *in vivo*. Doxy-treated or mock control mice were challenged I.P. with 1-3 x 10^8^ CFU of GAS strain NV19 (NV6 + pDC-sgRNA-*ftsZ*) and monitored for disease progression. Doxycycline (25 μg for ∼1 mg/kg) or vehicle control were administered I.P. 1h before infection and 48h post-infection. As shown in Fig. 5D, 80% of mice that received doxy survived after 14 days compared to only 30% of mock-treated mice (p = 0.0027, Mantel-Cox Log Rank test). To ensure that bacterial clearance was not solely attributed to the doxy treatment, an additional control group of mice were infected with strain NV26 (NV6 + pDC-sgRNA-*control*), carrying a non-targeting control sgRNA. No statistically significant difference in survival was observed for this group (**Fig. 5D**).

## Discussion

GAS remains a predominant cause of bacterial infections worldwide. While significant strides have been made in understanding GAS physiology and its interactions with the host (1, 17), fundamental insights into its basic cell biology lag behind. Such knowledge is crucial for the development of vaccines and the identification of novel drug targets. The existing knowledge gap is largely attributed to the challenges in genetic manipulation of pathogenic GAS strains and the absence of a modern molecular toolbox. Here, we bridge this gap by creating a highly transformable, fully virulent M1T1 GAS strain, establishing a titratable doxy/tetracycline/aTc-inducible system, and introducing CRISPRi for controlled gene expression in GAS.

By employing a non-polar *hsdR* mutant, sgRNAs can be directly cloned into GAS without the need for an *E. coli* intermediate. This streamlined approach makes it possible to obtain a specific gene depletion within days, as opposed to weeks or months (10). Utilizing readily available oligonucleotides, following the design outlined in Table S1, and using previously prepared electrocompetent NV6 cells (*hsdR::ery*, *cas9::tetM*, *tetR*, Ptet-*dcas9*) a CRISPRi experiment can be conducted in as little as three days. Our system would also be easily transferable to other GAS strains by the addition of the phage anti-restriction protein Ocr, which protects incoming DNA from restriction by the HsdR system, thus negating the need to work in an *hsdR* mutant background (47). Importantly, we show that not only is this CRISPRi system functional *in vitro*, but it can be efficiently induced in a murine infection model, simply by administering doxy to the animal. Many GAS strains pick up *covS* mutations during *in vivo* infections, rendering them hypervirulent (48). Therefore, future *in vivo* CRISPRi experiments might be more efficient when performed in preexisting *covS* mutant strains. We also note that CRISPRi works by repressing transcription and not at the translational level and due to its polar effects, genes downstream, or sometimes even upstream, might be affected when targeted by CRISPRi (31, 32, 49). Also, when targeting essential genes, *dcas9* mutants may be selected for, which needs to be taken into account when interpreting CRISPRi data (40).

The tools and methodologies outlined in this study can markedly enhance the throughput by which GAS research may be performed, and we foresee that genome-wide CRISPRi screens (CRISPRi-seq) will soon become feasible for GAS (40). CRISPRi and CRISPRi-seq offer advantages over existing TrAsh, Tn-seq and TRADIS workflows for GAS (11, 18, 50) since they enable the examination of essential gene functions and the performance of titratable drug-gene interaction studies. Moreover, we anticipate that the P*tet-dcas9* cassette and pDC-sgRNA plasmid system can be readily applied to other low GC-rich Gram-positive bacteria.

Annotation databases like SubtiWiki (51), EcoCyc (52), and PneumoBrowse (53) have significantly accelerated gene discovery, functional analysis, and hypothesis-driven studies for these bacteria. In this work, we introduce SpyBrowse (https://veeninglab.com/SpyBrowse), a public domain resource enabling users to explore the newly assembled GAS *S. pyogenes* 5448 NV1 genome. SpyBrowse facilitates the inspection of encoded features, regulatory elements, repeat regions, and other valuable properties. It also provides information on sgRNA binding sites and links to useful bioinformatic resources such as Alphafold predictions (54) (via Uniprot (14)) and PaperBlast (15). SpyBrowse will undergo regular updates and is fully capable of incorporating various omics data and improved genome annotations, akin to the approach taken by PneumoBrowse (53). The methodologies presented in this study can serve as a roadmap for developing CRISPRi in other challenging-to-transform bacteria. Together with SpyBrowse, they should represent a valuable resource for researchers in the GAS field.

## Methods

### Bacterial strains and culture conditions

All strains, plasmids and primers used are listed in Table 1 and Table S2. All GAS strains in this study are derivatives of *S. pyogenes* 5448 (12) and are listed in Table 1. Strains were grown in liquid THB (Hardy Diagnostics (or Oxoid THB for Fig. 1D, right panel) without aeration at 37°C. The IPTG-inducible promoter on pAV488 was activated with 1 mM IPTG (β-D-1-thiogalactopyranoside, Sigma-Aldrich). Electrocompetent GAS cells were made by overnight growth in THB + 0.6% glycine followed by 1:10 dilution in fresh THB + 0.6% glycine. If required, 1 mM of IPTG was used to induce recombineering from pAV488 (Varble, laboratory collection) when cells reached an OD600 of 0.15. At OD600 of 0.3, cells were centrifuged (4000 rpm, 10 min, 4°C) and washed four times with 0.625 M Sucrose before resuspending in 20% glycerol and stored at −80°C in 50 µl aliquots. For electroporation, DNA was added to one 50 µl aliquot of electrocompetent cells on ice for 5 min before transferring to a 1 mm electroporation cuvette (Genesee Scientific). After electroporation (Eppendorf 2510, 1.7 kilovolts), cells were incubated in THB + 0.25 M Sucrose for 2 h before overnight antibiotic selection on THA (37°C). When appropriate, the medium was supplemented with the following antibiotics: chloramphenicol (2 µg.mL^-1^), erythromycin (0.5 µg.mL^-1^), kanamycin (400 µg.mL^-1^), and tetracycline (0.5 µg.mL^-1^). Genomic GAS DNA was prepared using the Zymo Quick-DNA Fungal/Bacterial Miniprep Kit (4 ml of overnight culture was used as input). We note that some of the GAS strains engineered here contain an erythromycin resistance cassette. Since macrolide antibiotics are sometimes prescribed to treat GAS infections in individuals that have a penicillin allergy (all GAS strains described here are fully penicillin susceptible), we urge the community to only use these strains within appropriate biosafety conditions. We also note that in the unlikely event that someone with a penicillin allergy gets a GAS infection caused by one of these erythromycin-resistant strains, they can be safely treated with cephalexin or cefadroxil as all reported strains here are fully susceptible to these antibiotics.

Plasmid pDC-sgRNA was made in *E. coli* strain MC1061 (MClab) grown in LB medium at 37°C with aeration; 5 µg.mL^-1^ chloramphenicol was added when appropriate.

### Recombineering, plasmid and strain construction

Recombineering plasmid pAV488 (Varble, laboratory collection) was transformed by electroporation to wildtype 5448 (strain NV1) selecting on chloramphenicol resulting in strain NV2 (Table 1). Next, the recombineering enzymes Gam, ERF recombinase and single stranded DNA binding protein encoded on pAV488 were induced with 1 mM of IPTG and cells were made electrocompetent and stored at −80°C. To generate an *hsdR* replacement mutant, an erythromycin cassette without promoter and terminator but with its own RBS was amplified by PCR using primers ONV31/ONV32 (Table S2) using chromosomal DNA of strain VL4321 (55) as template. The *hsdR* upstream and downstream regions were amplified using chromosomal DNA of strain NV1 as template using primers ONV29/ONV30 and ONV33/ONV34, respectively (**Fig. 1**). The three PCR fragments were purified (Zymo DNA Clean and Concentrator kit: ‘Zymo kit’) and used in a 1:1:1 molar ratio Golden Gate assembly reaction with BsaI (NEB) and T4 ligase (NEB) for 50 cycles of 1.5 min at 37°C followed by 3 min at 16°C. Enzymes were inactivated at 80°C for 10 min. The assembly was purified (Zymo kit) and 1 µl was used as a template in a PCR using primers ONV29/ONV34. The resulting 3431 bps fragment was purified (Zymo kit) and 10 µl was used to transform electrocompetent NV2 cells. Erythromycin resistant colonies were selected and used for further analysis resulting in strain NV3 (*hsdR::ery*, pAV488). Strain NV3 was cured from plasmid pAV488 by growing it overnight in THB with 1 mM IPTG (without chloramphenicol) and restreaking single colonies, resulting in strain NV28 (*hsdR::ery*).

Strain NV4 (*hsdR::ery*, *cas9::kan*, pAV488) was made by amplifying approximately 1kb upstream and downstream of the *cas9* locus (**Fig. 2B**) using primers ONV1/ONV2 and ONV5/OV6, respectively, using chromosomal DNA of strain NV1 as template. A kanamycin resistance cassette including promoter and terminator was amplified from plasmid pAV258 (Varble laboratory collection) using primers ONV3/ONV4. The three PCR fragments were purified (Zymo kit) and used in a 1:1:1 molar ratio Golden Gate assembly reaction with AarI (ThermoFisher) and T4 ligase (NEB) for 50 cycles of 1.5 min at 37°C followed by 3 min at 16°C. Enzymes were inactivated at 80°C for 10 min. The assembly was purified (Zymo kit) and 1 µl was used as a template in a PCR using primers ONV1/ONV6. The resulting 3995 bps fragment was purified (Zymo kit) and 10 µl was used to transform electrocompetent NV3 cells. Kanamycin resistant colonies were selected and used for further analysis resulting in strain NV4.

Strain NV6 (*hsdR::ery*, *cas9::tetM, tetR,* Ptet*-dcas9*) was constructed by amplifying approximately 1 kb upstream and downstream of the *cas9* locus using primers ONV1/ONV9 and ONV6/ONV12, respectively, using chromosomal DNA of strain NV1 as template. The *tetM-tetR-Ptet-dcas9* cassette was amplified from chromosomal DNA of strain VL3469 (36) using primers ONV10/ONV11. The three PCR fragments were purified (Zymo kit) and used in a 1:1:1 molar ratio Golden Gate assembly reaction with AarI (ThermoFisher) and T4 ligase (NEB) for 60 cycles of 1.5 min at 37°C followed by 3 min at 16°C. Enzymes were inactivated at 80°C for 10 min. The 9774 bps ligation product was cut from gel and purified (Zymoclean Gel DNA recovery kit) and directly used to transform electrocompetent NV4 cells. Tetracycline resistant colonies were selected and plasmid cured, resulting in strain NV6. Since NV6 cells harbor the *tetM* resistance marker, growth is not perturbed by doxy upon inducing with 20 ng/ml.

Plasmid pDC-sgRNA was constructed as follows: A PCR using primers ONV21/ONV22 and plasmid pDC123 (24) as template was performed to obtain the vector backbone. The *sgRNA-mCherry* cassette flanked by the Illumina read 1 and read 2 sequences were amplified using plasmid pVL4930 (Veening laboratory collection) as template with primers ONV23/ONV24. Next, the two fragments were purified (Zymo kit) and used in a 1:1 molar ratio Golden Gate assembly reaction with BsaI-HF2 (NEB) and T4 ligase (NEB) for 50 cycles of 1.5 min at 37°C followed by 3 min at 16°C. Enzymes were inactivated at 80°C for 10 min. The ligation product was used to transform chemically competent *E. coli* MC1061 (MCLab). A pink, mCherry-expressing chloramphenicol resistant colony was restreaked and selected for further analysis resulting in *E. coli* strain pDC-sgRNA (Table 1). Plasmid pDC-sgRNA was purified (Qiagen miniprep kit) and verified by nanopore sequencing (Plasmidsaurus). Note that *mCherry*, and the subsequent sgRNA, is driven by the strong constitutive P3 promoter (42) and that all cloned sgRNAs will have adenine as initiating nucleotide (+1) (**Fig. 2A**), ensuring strong expression regardless of spacer sequence (40, 53).

To construct GAS strains NV19, NV22, NV25 and NV26, complementary oligos ONV60/ONV61, ONV66/ONV67, ONV72/ONV73 and ONV74/ONV75, respectively, were annealed and phosphorylated. Briefly, 2.5 µl of each complementary oligo (at 100 µM concentration) were annealed in a 50 µl reaction containing 5 µl 10x TEN buffer (100 mM Tris-HCl pH8, 10 mM EDTA, 500 mM NaCl) for 5 min at 95°Ϲ and slowly cooled to room temperature. Next, 1 µl of the annealed oligos were phosphorylated in a 10 µl reaction containing 0.25 µl T4 PNK (10000 units/ml, NEB) and 1 µl 10x T4 ligase buffer for 40 min at 37°Ϲ before heat-inactivation at 65°Ϲ for 20 min. Finally, the phosphorylated annealed oligos were diluted 10-fold (to 0.05 µM DNA) ready to use for ligation. To generate the digested pDC-sgRNA vector, 1 µl of plasmid pDC-sgRNA was used as a template in a PCR using outward facing primers ONV76/ONV77 binding within the *mCherry* sequence. The PCR product was incubated with EcoRV (NEB) that cuts inside *mCherry* for 30 min at 37°C to linearize any remaining template DNA. Next, the reaction was purified (Zymo kit) and digested with BsmBI-HF (NEB) at 55°C for 3h followed by purification using the Zymo kit. 100 ng of pure digested pDC-sgRNA was used in a 15 µl ligation reaction containing 3.5 µl of the 0.05 µM annealed phosphorylated primers. Ligation was performed for 1h at RT or overnight at 16°C. Finally, 7 µl of the ligation mixture was used to transform electrocompetent cells of strain NV6 by electroporation, resulting in strains NV19, NV22, NV25 and NV26, respectively. Plasmids were isolated by miniprep (Qiagen) and correct spacer sequences were verified by Sanger sequencing using primer ONV92.

### Microtiter plate-based growth assay

Overnight GAS cultures grown in THB medium at 37°C were diluted 1:20 in the morning in fresh THB until mid-exponential growth (OD_600nm_ = 0.3) with no inducer at 37°C, after which they were diluted to OD_600nm_ = 0.004 in 250ul of fresh THB medium supplemented with doxy when appropriate inside 96-well flat bottom microtiter plates (Costar 3370) covered with Breath-Easy film (Sigma) to prevent evaporation. Cellular growth was then monitored every 10 min at either 37°C in a microtiter plate reader (TECAN Infinite F200 Pro). Each growth assay was performed in triplicate and the average of the triplicate values with standard errors of the mean were plotted using BactExtract (56).

### DIC and fluorescence microscopy

GAS cells were grown in THB medium at 37°C to an OD_600nm_ = 0.3 without any inducer and diluted 50 times in fresh THB medium supplemented, when appropriate, with 20 ng.mL^-1^ doxy (for activation of dCas9). After 3 h, 1 mL of culture was collected. For membrane staining, 5 µg.mL^-1^ of Nile red (Invitrogen) was added and for DAPI staining, 1 µg.mL^-1^ DAPI (Sigma-Aldrich) was added to the cells and incubated for 5 min at room temperature prior to centrifugation. Cells were washed twice with 1 mL PBS and re-suspended into 50 µL PBS. 1 µL of cells were then spotted onto PBS agarose (1%) pads in 10-well multitest microscope slides (MP Biomedicals). Microscopy acquisition was performed using a Zeiss M1 upright microscope with a 100x oil-immersion objective. Images were processed using Fuji (57).

### Genome sequencing, annotation, methylation analysis and spacer design

Strain NV1 (GAS 5448, Nizet lab collection) was sequenced, assembled, and annotated by Plasmidsaurus, Inc. according to the protocols listed on the Plasmidsaurus website (https://www.plasmidsaurus.com/faq/#bact-assembly). In addition, strains NV1, NV28 and NV6 were also sequenced using the PacBio Sequel II instrument at the Lausanne Genomics Facility. Read demultiplexing and quality control were performed with SMRTLink version 11.0 (https://www.pacb.com/support/software-downloads/). The microbial genome assembly pipeline in this toolkit was used to assemble genomes and identify methylation motifs. Assemblies were circularized with circlator (58). Structural variants were detected with pbmm2 version 1.13.1 (59) and pbsv version 2.9.0 using default settings then confirmed manually. Whole genome alignments were performed and visualized in Mauve (60). This revealed a large genome inversion of 1,475,033 base pairs. Prokka (43) was used to annotate the NV1 PacBio genome and used to design a unique sgRNA (see Table S1) for every genetic feature as described (40). Genome sequences confirmed correct gene replacement of *hsdR* in NV6 and NV28 and correct integration of Ptet-*dcas9* at the *cas9* locus in NV6. Besides the large genome inversion and several smaller insertions, the genome differs on the following sites from the publicly available 5448 genome sequence (25) (GenBank: CP008776.1): NV1 numbering: 196’187 t-->c, 196’188 c-->t, 196’385 g-->t, 196’446 t-->c, 196’452 t-->c, 296’177 a-->c, 296’508 g-->t, 296’568 t-->c, 296’575 t-->c, 334’494 c-->a, 334’434 a-->g, 334’427 a-->g, 1’308’897 g-->a, 1’310’949 a-->c, 1’518’946 g-->a, 1’518’953 g-->a, 1’519’013 a-->c, 1’565’892 a-->t, 1’599’979 a-->c, 1’672’223 a-->g, 1’672’224 g-->a). These SNPs are mainly related to the ISAs1-like elements. The polished NV1 genome was further annotated by the NCBI Prokaryotic Genome Annotation Pipeline (PGAP) and is available with accession number CP140117.2 and visualized through SpyBrowse (see below).

### SpyBrowse

SpyBrowse (https://veeninglab.com/SpyBrowse) is based on JBrowse 2 (13). Features were divided over 3 annotation tracks: (i) Reference sequence, (ii) Coding features, and (iii) designed sgRNAs. GC content is calculated via the NucContent plugin (available via: https://github.com/jjrozewicki/jbrowse2-plugin-nuccontent). The NCBI annotation of *S. pyogenes* NV1 was used for the coding features track (accession: CP140117.2)

### Quantifications and statistical analysis

Data analysis was performed using R (R version 4.2.2) and Prism (Version 10.0.3, Graphpad). Data shown are represented as the mean of at least three replicates ± SEM if data came from one experiment with replicated measurement, and ± SD if data came from separate experiments.

### Flow cytometry

NV25 (sgRNA-*emm*), NV26 (sgRNA-control), and M protein knockout GAS M1T1 5448 were grown overnight in THB with chloramphenicol for NV25 and NV26. Cultures were diluted 1:10 from overnights in THB (without chloramphenicol) and grown to mid-logarithmic phase (OD_600_ = 0.4) for 2.5 hours. During this time, NV25 and NV26 were grown +/-20 ng/ml doxycycline to induce the CRISPRi system. Bacteria were washed in PBS and incubated in 10% donkey serum at room temperature. Rabbit M protein antisera (Vaxcyte) was added at 2% final concentration for 1 hour at room temperature. Bacteria were washed with PBS and then incubated in 1:200 donkey anti-rabbit IgG conjugated AlexaFluor 488 fluorophore (Thermo Fisher #21206) for 30 minutes at room temperature. Samples were washed in PBS and run on a BD FACSCalibur. Flow cytometry data were analyzed with FlowJo v. 10.8.2 (Tree Star, Inc.).

### Mouse infection experiments

Eight-week old female CD1 mice (Charles River Laboratories) were infected intraperitoneally with 1-3 x 10^8^ CFU in 100 μl of each GAS engineered strain (n = 20 per strain). One hour prior to infection, mice were injected intraperitoneally with either 25 μg doxycycline (n = 10) or PBS (n = 10). 48h following infection a second 25 μg doxy dose was administered i.p. Mortality was observed daily for 14 days post infection. Mice were housed on a 12 hour light/dark schedule, fed a 2020x diet (Envigo), and received acidified water. Prior to experimentation, mice were randomized into cages with no more than 5 mice per cage. Mouse experiments were approved by the UC San Diego Institutional Animal Care and Use Committee (Protocol #S00227M) in compliance with federal regulations and were conducted with oversight from veterinary professionals.

## Data availability

The data that support the findings of this study are incorporated in the manuscript and its supporting information. Pacbio genome sequences, assemblies, and sequencing reads are available at NCBI under BioProject accession number PRJNA1050251.

## Acknowledgements

We thank the members of the Nizet and Veening groups for valuable discussions. We thank Doran Pauka and Colin Diesh for help with SpyBrowse. We thank the Lausanne Genomic Technologies Facility for SMRT sequencing and continued support. J.W.V. received financial support to carry out this research at UCSD through the Fondation Herbette and by the Swiss National Science Foundation (SNSF) (Scientific Exchange grant IZSEZ0_213879). A.B.J. is supported through a Postdoctoral Fellowship grant (TMPFP3_210202) by the SNSF. Work in the lab of J.W.V. was supported by SNSF grants 310030_192517, 310030_200792 and 51NF40_180541.

## Author contributions

J.W.V. wrote the paper with input from all authors. J.W.V., E.B., A.S., A.B.J., P.S.G., S.D., J.L.B. and A.M.B. performed the experiments. J.W.V. and V.N. designed, analyzed and interpreted the data.

## Supplementary information

Supplementary Table S1: GAS 5448 (NV1) genome-wide sgRNA design

Supplementary Table S2: Oligonucleotides

